# Accelerating CHO-K1 Cell Line Development by Reducing Suspension Adaptation with a Microplate Agitation Culture System

**DOI:** 10.64898/2025.12.11.693844

**Authors:** Shih-Pei Lin, Ching-Nan Lin, Wei-Rou Wang, Cheng-Han Tsai

## Abstract

Stable and productive CHO cell lines are essential for biopharmaceutical manufacturing, yet early expansion steps are often constrained by prolonged period required for suspension adaptation. Single-cell cloning (SCC) ensures monoclonality and regulatory compliance, but cells transitioning from static to suspension culture frequently exhibit variable recovery, which prolongs timelines and increases process variability. To address this challenge, mixing-based microplate culture systems have been developed to improve early expansion efficiency. The C.NEST platform provides controlled pneumatic mixing and environmental monitoring that facilitates earlier adaptation to suspension conditions. At the 96-well and 24-well stages, this system allows cells to establish stable growth under suspension-like environments, thereby shortening the adaptation period following transfer to shaking culture. In this study, we applied C.NEST to the SCC workflow for developing CHO-K1 stable cell lines. Integrating C.NEST’s controlled mixing reduced adaptation time, enhanced the consistency of clone expansion, and improved the ability to identify high-yield clones. These findings highlight the potential of C.NEST to streamline cell line development workflows by accelerating early suspension adaptation and improving clone selection reliability.

**Graphical abstract:** 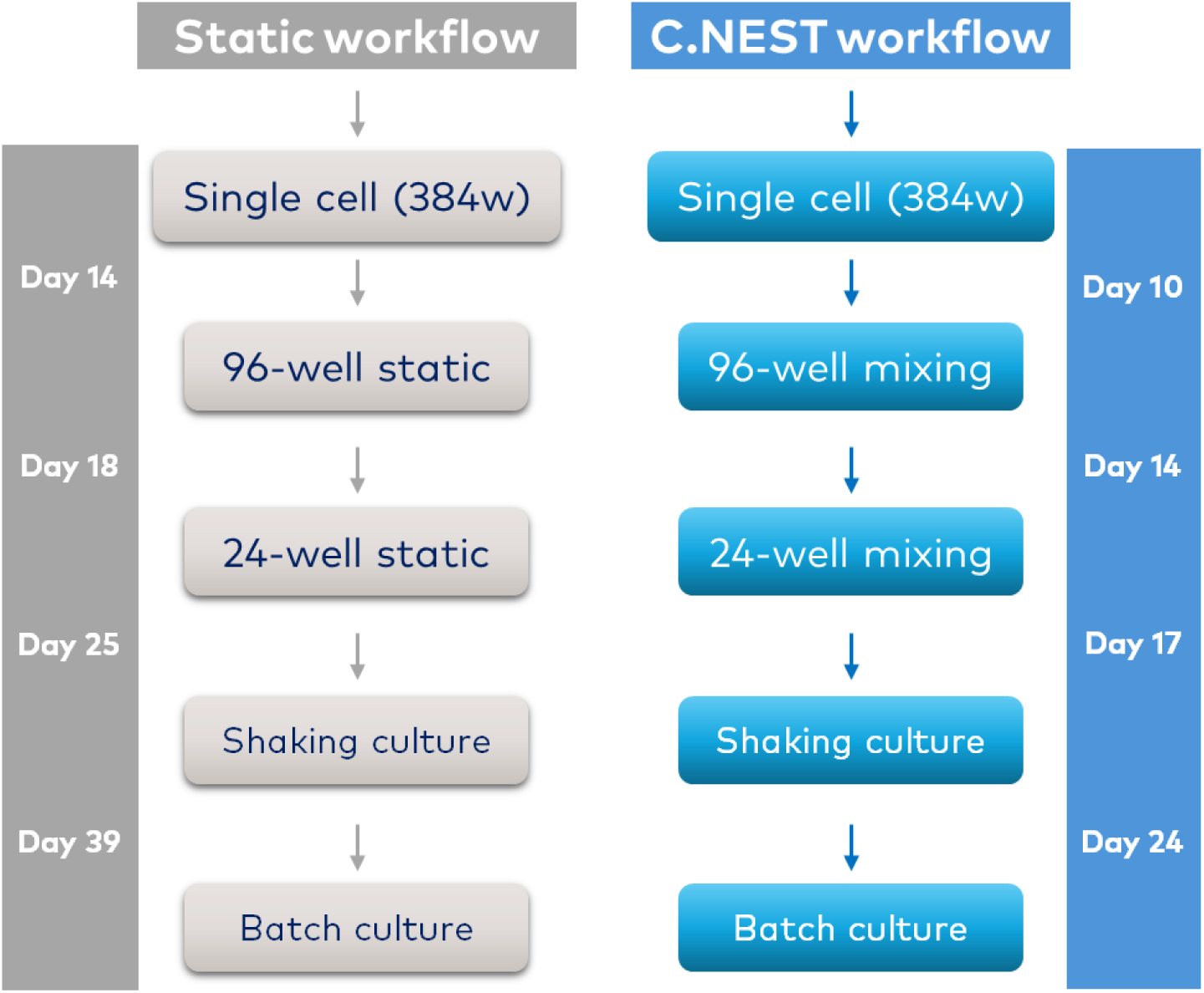

C.NEST mixing shortens suspension adaptation, accelerates clone expansion, and enhances early-stage screening.

**Highlight:**

- The C.NEST microplate agitation culture system accelerates early CHO-K1 cell line development.
- Controlled pneumatic mixing improved oxygen transfer and medium homogeneity, promoting stable growth during early expansion.
- Early mixing shortens suspension adaptation by approximately one week.
- Mixing cultures enabled more accurate clone performance assessment, revealing high-producing outliers.
- C.NEST provides a scalable and reproducible solution for integrating mixing-based culture into single-cell cloning workflows.

## Introduction

The development of stable and high-yield Chinese hamster ovary (CHO) cell lines remains a critical step in biopharmaceutical manufacturing, with more than 70% of recombinant therapeutic proteins currently produced in CHO-based systems (Walsh, 2022 [1]; Kim et al., 2012 [2]; Jayapal et al., 2007 [3]). Traditionally, SCC has been the cornerstone of cell line development (CLD), enabling the isolation of monoclonal populations to ensure consistency in growth, productivity, and product quality (ICH Q5D, 1998 [4]; EMA, 2006 [5]; FDA, 1992 [6]).

During the early stages of clone expansion, cells must transition from static culture to suspension growth, a process known as suspension adaptation, which substantially affects growth kinetics and overall clone performance (Wurm, 2013 [7]; Fischer et al., 2015 [8]). This adaptation phase frequently represents a bottleneck in CLD, prolonging development timelines and increasing variability.

To overcome these limitations, microplate agitation culture systems such as the C.NEST (Leadgene Biosolutions (formerly CYTENA BPS)) platform have been introduced. Originally developed to support clonal expansion from single cells, C.NEST provides controlled mixing and environmental monitoring that facilitates earlier adaptation to suspension culture. At the 96-well and 24-well stages, the system enables cells to establish suspension growth earlier in the workflow, thereby accelerating proliferation and reducing the adaptation period required after transfer into shaking culture.

Unlike mechanical or platform-based agitation methods, the pneumatic-driven mixing in C.NEST is generated through periodic pressure modulation within a closed chamber. By alternating controlled pulses of gas inflow and outflow, the system induces a gentle vertical recirculation of the culture medium. This mode of media movement minimizes shearing forces while improving convective transport across the well depth, promoting more uniform distribution of nutrients, metabolites, and dissolved gases. As a result, local concentration gradients, commonly observed in static cultures or insufficiently mixed microplate systems, are mitigated.

A notable advantage of this approach is that gas exchange and mixing occur simultaneously at the gas-liquid interface, providing a more efficient pathway for oxygen dissolution compared with orbital mini-shakers, which rely on surface agitation and often generate uneven gas transfer across wells. Because the plate remains stationary, the system avoids issues such as micro-bubbles, edge-well evaporation, splashing, and cross-contamination commonly associated with mechanical movement of microplates. The ability to independently set mixing parameters per chamber further allows fine-tuning of oxygen transfer rates and mixing intensities to match cell type or stage-specific requirements. This degree of control is rarely achievable in conventional early-stage culture formats.

In this study, we evaluated the application of C.NEST mixing to accelerate clone expansion and reduce suspension adaptation time in CHO-K1 cells. Our goal was to streamline the CLD workflow and improve the consistency and sensitivity of early-stage clone performance assessment.

## Material and method

### Cell Culture

Suspension-type CHO-K1 and mAb-expressing CHO-K1Q cell lines (QuaCell Biotechnology) were used in this study. mAb-CHO-K1 cells were maintained in 125 mL shake flasks (Thermo Fisher Scientific, #4115-0125) containing Dynamis medium (Gibco, #A2661501) supplemented with 4 mM L-glutamine (Corning, #25-005-CI) and penicillin–streptomycin (Corning, #30-0020-CI), with a working volume of 30 mL. Cultures were incubated at 37 °C, 5% CO₂, and 130 rpm on a 19 mm orbital shaker. mAb-CHO-K1Q cells were maintained in EX-CELL® CD CHO Fusion medium (Merck, #24365C) under the same incubation conditions. Cells were passaged every 3-4 days and reseeded at 3 × 10⁵ cells/mL. Cell density and viability were measured using trypan blue exclusion and quantified on a FACSCOPE B Cell Counter (Curiosis).

### Microplate Agitation Culture in C.NEST

mAb-CHO-K1 cells were cultured in 96-well (Corning, #3599) and 24-well (Greiner Bio-One, #662102) tissue culture plates under static or mixing conditions. For the 96-well format, cells were seeded at 0.5-5 × 10⁵ cells/mL in 200 µL per well and cultured for 4 days. For the 24-well format, cells were seeded at 0.2-10 × 10⁵ cells/mL in 1.4 mL per well and cultured for 3 days. Static groups were incubated under standard conditions (37 °C, 5% CO₂), while mixing groups were cultured on the C.NEST platform with cycle settings of 50 s (C.NEST_50s, 96-well) or 10 s (C.NEST_10s, 24-well). No feeding was applied during these short-term cultures. At the defined endpoints (Day 4 for 96-well, Day 3 for 24-well), viable cell density was determined by trypan blue exclusion and quantified with a FACSCOPE B Cell Counter. Statistical analyses were performed using two-way ANOVA. Significance thresholds were set as follows: ns, P > 0.05; *P < 0.05; **P < 0.01; ***P < 0.001; ****P < 0.0001.

### Single-Cell Seeding and Expansion Workflow

Single-cell seeding was performed using the UP.SIGHT system (CYTENA) with automated dispensing and monoclonality confirmation. Cells were deposited into 384-well plates (Corning, #3680) with 50 µL cloning medium per well, consisting of 80% EX-CELL® CHO Cloning Medium (Merck, pre-supplemented with 4 mM L-glutamine) and 20% conditioned medium. Plates were incubated at 37 °C and 5% CO₂ under static conditions and monitored for outgrowth using the integrated imaging system.

Wells with >40% outgrowth were considered for expansion. Confluency was evaluated using UP.SIGHT or the CloneSelect Imager (Molecular Devices), with a threshold of >10%. For one-to-two well splitting experiments, transfers were only performed once confluency reached ≥20%, ensuring sufficient viable cell numbers for further growth.

Selected clones were transferred on Day 10 into 96-well plates (200 µL per well) and subsequently into 24-well plates (1.4 mL per well). Both formats were cultured on the C.NEST platform (50 s/cycle in 96-well, 10 s/cycle in 24-well). Expansion continued into TubeSpin® Bioreactor tubes (TPP, #87050; 10 mL, 230 rpm) or 125 mL shake flasks (20 mL, 130 rpm, 19 mm orbit). Cultures were maintained at 37 °C, 5% CO₂. At each expansion step, viable cell density and viability were assessed by trypan blue exclusion and quantified on a FACSCOPE B Cell Counter.

### IgG Titer Estimation Using F.QUANT

Relative IgG titers were determined using the F.QUANT Fc Titer Assay (CYTENA), a bead- and plate-based fluorescence assay. Supernatants were clarified and loaded onto Fc-specific assay plates according to the manufacturer’s instructions. Fluorescence intensity was used to rank clones, and top producers were advanced for further expansion and batch culture evaluation.

## Result

### Enhanced Cell Expansion by C.NEST Mixing in Microplate Cultures

To evaluate the effect of C.NEST mixing on early-stage cell expansion, we compared viable cell density (VCD) in 96-well and 24-well plates under mixing conditions versus static culture. After 4 days of cultivation in 96-well plates, clones cultured with 50 s mixing (C.NEST_50s) exhibited significantly higher VCD compared to static culture, particularly at seeding densities ≥1 × 10⁵ cells/mL (Figure 1A). Similarly, after 3 days of culture in 24-well plates, cells exposed to 10-second mixing (C.NEST_10s) demonstrated markedly improved growth over static controls, with enhanced VCD observed at initial densities above 0.5 × 10⁵ cells/mL (Figure 1B). These results indicate that the pneumatic-driven mixing implemented by C.NEST accelerates suspension adaptation and supports more robust cell expansion across both 96- and 24-well formats.

**Figure 1.**
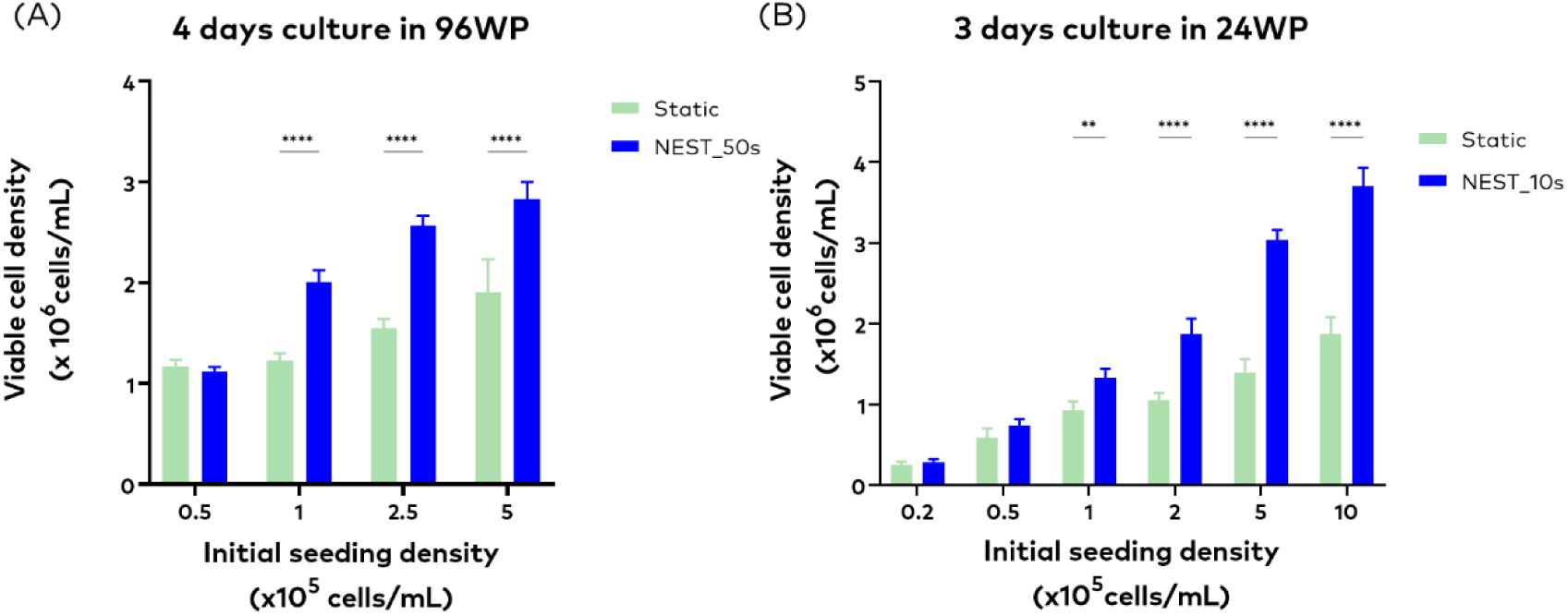
Effect of NEST mixing culture on cell growth in 96- and 24-well plates. Viable cell density measured after (A) 4 days of culture in 96-well plates with 50 s mixing (NEST_50s) or static conditions, and (B) 3 days of culture in 24-well plates with 10 s mixing (NEST_10s) or static conditions. NEST mixing significantly enhanced cell growth at initial seeding densities above 1 × 10⁵ cells/mL in 96-well plates and above 0.5 × 10⁵ cells/mL in 24-well plates. Data are mean ± SD; P < 0.01 (**), P < 0.0001 (****).

### Defining Confluency Thresholds for Initiating 96-Well Mixing

To establish the criteria for initiating mixing culture, we examined the relationship between confluency and viable cell numbers (Figure 2). Bright-field images illustrated four representative confluency levels: Level 1 (<10%), Level 2 (10–40%), Level 3 (40–70%), and Level 4 (>70%) (Figure 2A). Subsequent quantification of viable cell counts confirmed that wells exceeding 10% confluency typically contained close to (approximately) 1 × 10⁴ viable cells. As cell numbers substantially increased, the Image system classified these cultures into higher confluency levels: Level 3 and Level 4 (Figure 2B).

**Figure 2.**
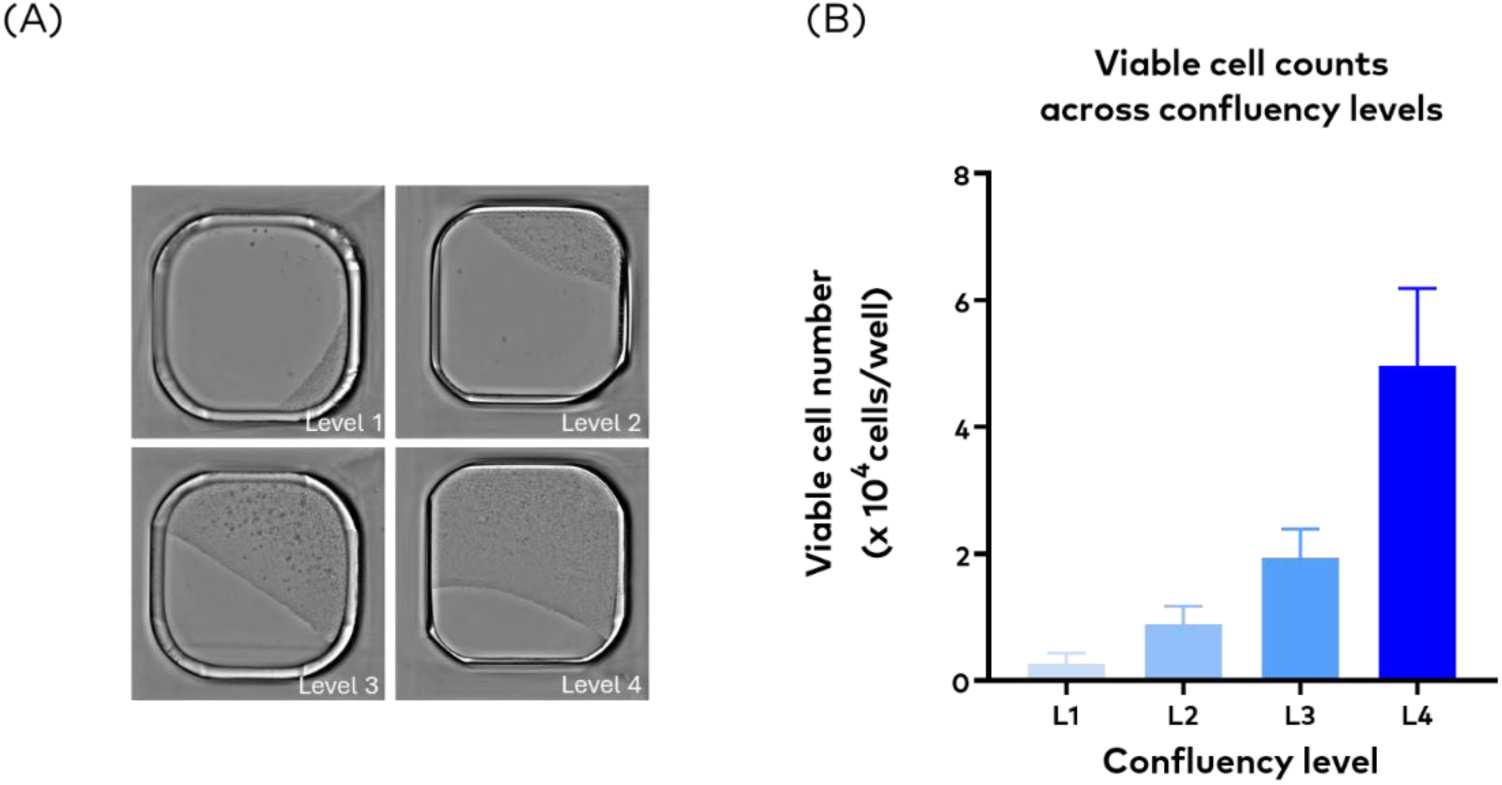
Relationship between cell confluency and viable cell number. (A) Representative bright-field images illustrating four confluency levels: Level 1 (<10%), Level 2 (10–40%), Level 3 (40–70%), and Level 4 (>70%).(B) Viable cell counts (×10⁴ cells/well) corresponding to each confluency level. Wells exceeding 10% confluency typically contained ∼1 × 10⁴ viable cells (200 µL per well, equivalent to 5 × 10⁴ cells/mL), which was sufficient to initiate 96-well mixing culture. This threshold served as the general criterion for determining suitability of wells for mixing. In validation experiments requiring synchronized parallel workflows, a stricter threshold (∼30% confluency) was applied to account for splitting clones from the same 384-well origin and potential cell loss during transfer. Data represent mean ± SD from n =7.

Previous tests indicated that premature mixing initiation or starting with low cell densities failed to accelerate growth and was actually detrimental to cell expansion. Conversely, prolonging static culture to accumulate higher cell numbers compromised cell health and subsequently inhibited growth performance. Therefore, we defined the minimum requirement for reliable transfer into 96-well mixing as 1 × 10⁴ cells per well (200 µL culture volume, equivalent to a seeding density of 5 × 10⁴ cells/mL). This criterion ensured that clones could adapt efficiently and expand robustly under mixing conditions.

### Case 1: Validation of C.NEST mixing for early clone expansion

In the first validation case, paired clones derived from the same single-cell seeding (pool A) were expanded in parallel under static or C.NEST mixing conditions to directly compare workflow design (Figure 3). This schematic illustrates the two workflows: (1) the “Traditional” static expansion through 96-well and 24-well static plates before entering TubeSpin, and (2) the experimental C.NEST expansion under mixing conditions in both 96-well and 24-well stages prior to TubeSpin transfer. To synchronize the two workflows, clones were split from the same 384-well origin; moreover, a stricter threshold of ∼30% confluency was applied for initiating 96-well culture to account for potential cell loss during transfer.

**Figure 3.**
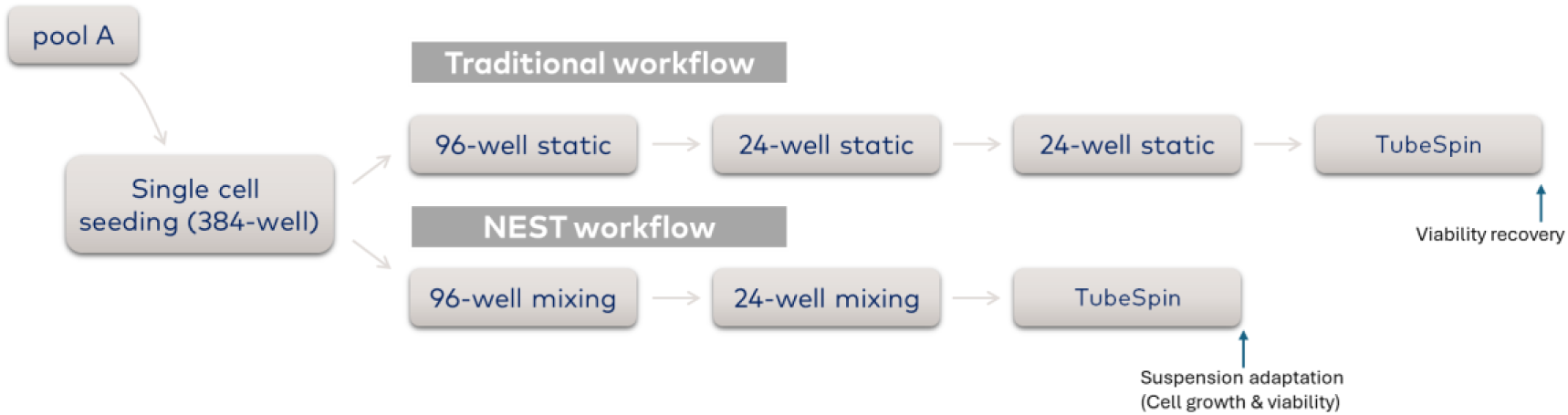
Case 1 schematic comparison of static and C.NEST mixing workflows. Clones derived from the same 384-well origin were split into parallel workflows. In the static workflow, clones progressed through 96-well and 24-well static culture before entering TubeSpin. In the C.NEST workflow, clones were expanded under mixing conditions in 96-well and 24-well plates prior to TubeSpin transfer. For synchronized parallel testing, a stricter threshold of ∼30% confluency was applied for initiating 96-well culture, ensuring sufficient viable cells were available for both workflows.

Subsequent monitoring of clone expansion demonstrated a clear advantage for the C.NEST mixing culture: clones exhibited significantly higher viable cell numbers after transferring to 24-well plates (Day 19, Figure 4B), despite comparable VCDs during the preceding 96-well stage (Day 16, Figure 4A). By day 23, C.NEST-derived clones had already progressed into TubeSpin culture, whereas static-derived clones remained restricted to 24-well plate, resulting in an evident divergence in expansion profiles (Figure 4C).

**Figure 4.**
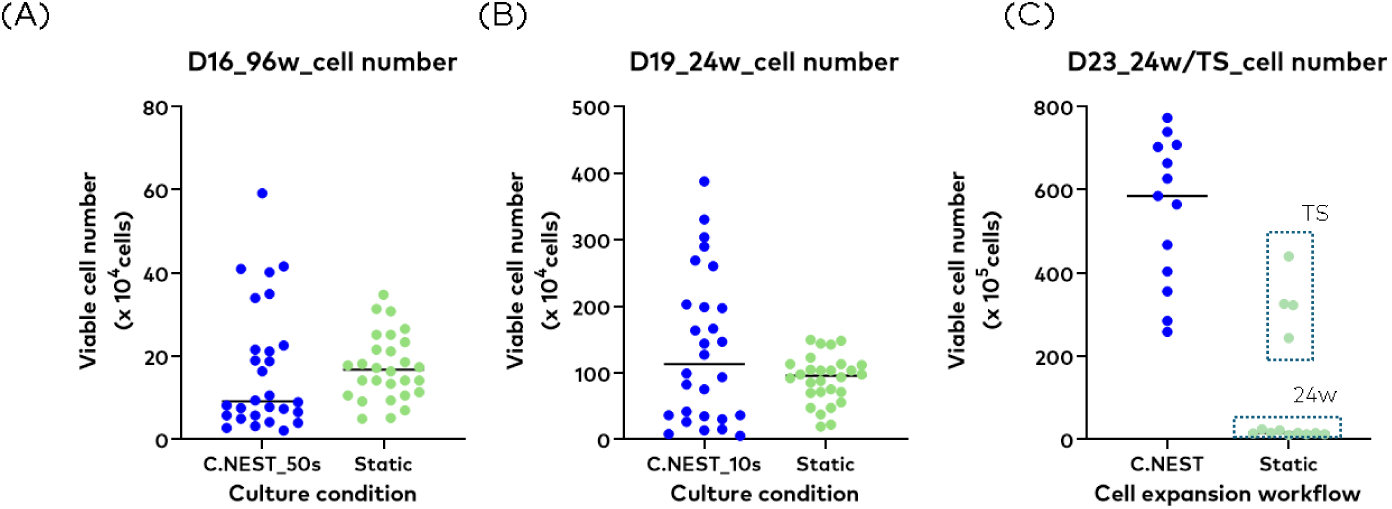
Case 1 –Effect of C.NEST mixing on clone expansion and progression to TubeSpin (TS) culture. (A) Viable cell numbers at day 16 after 4 days in 96-well plates under static or C.NEST_50s mixing conditions.(B) Viable cell numbers at day 19 after 3 days in 24-well plates under static or C.NEST_10s mixing conditions.(C) Viable cell numbers at day 23, where clones in the C.NEST workflow had advanced to the TubeSpin stage, whereas static clones remained in the 24-well stage. Mixing cultures displayed significantly higher viable cell counts compared to static controls.

On day 26, all clones were passaged into TubeSpin culture at 3 × 10⁵ cells/mL. Clones derived from the C.NEST workflow maintained higher viable cell numbers than those from the static workflow (Figure 5A). However, However, due to the relatively low productivity of the original pool, the average IgG titers of C.NEST-derived clones were only slightly elevated compared to static clones and did not reach statistical significance and did not reach statistical significance (Figure 5B). Therefore, a higher-producing pool needed to be selected to effectively validate the C.NEST workflow.

**Figure 5.**
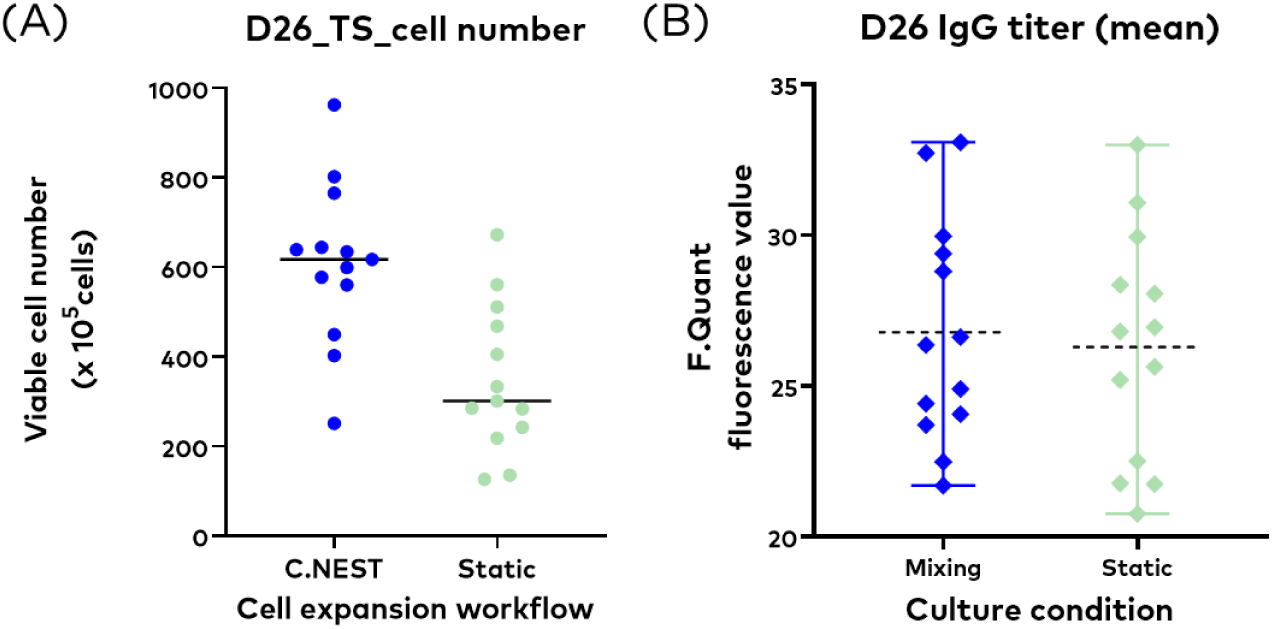
Case 1 –Clone performance at day 26 after passage at 3 × 10⁵ cells/mL. (A) By day 26, all clones could be passaged at a density of 3 × 10⁵ cells/mL. After 3 days of Tubespin culture, clones pre-expanded under C.NEST mixing maintained higher viable cell numbers compared to static-derived clones. (B) Because the original pool titer was too low, the average titer of mixing-derived clones appeared slightly higher than that of static-derived clones, but the difference was not significant. Therefore, a higher-producing pool was selected for further testing in the next stage.

The timeline (Figure 6) with more detailed cell passage criteria illustrates that the C.NEST mixing workflow enhanced cell growth, allowing clones to advance to TubeSpin culture earlier. Static-derived clones, however, required a longer incubation time in 24-well plates to reach the necessary VCD for suspension transfer. Together, these results demonstrate that C.NEST mixing accelerates early-stage clone expansion, although the low productivity of the original pool constrained the ability to discriminate clone performance. Together, these results demonstrate that C.NEST mixing accelerates early-stage clone expansion, although the low productivity of the original pool constrained the ability to discriminate clone performance.

**Figure 6.**
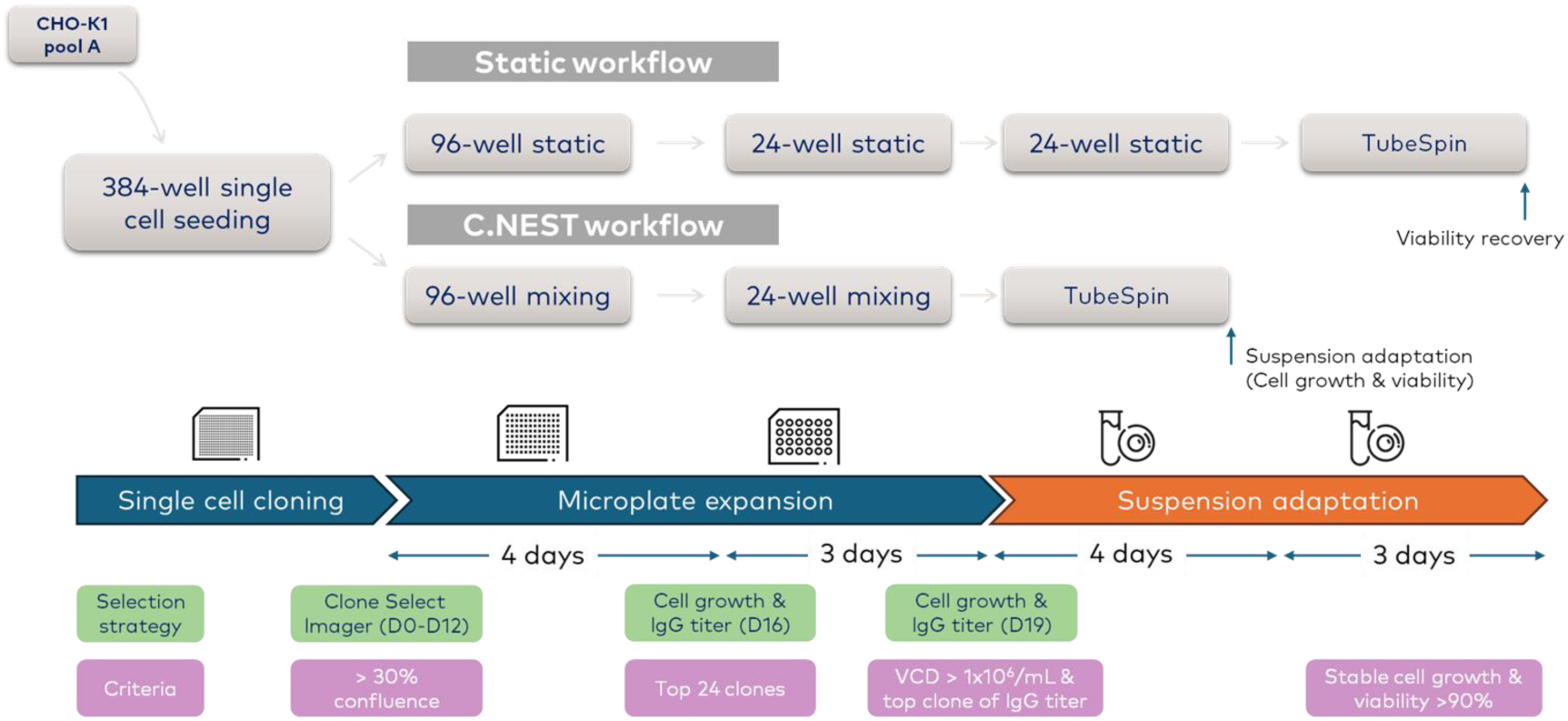
Case 1 workflow timeline for static and C.NEST mixing conditions. Paired clones derived from the same single-cell pool (pool A) were expanded under either static or C.NEST mixing workflows. In the C.NEST workflow, clones progressed through 96-well and 24-well mixing culture and advanced into TubeSpin suspension culture earlier. In contrast, static-derived clones required prolonged culture in 24-well plates to accumulate sufficient cell numbers before entering TubeSpin. The timeline (D14–D26) illustrates the duration of culture at each stage, highlighting the accelerated progression achieved with C.NEST mixing.

### Case 2: Evaluation of C.NEST mixing in clone selection and TubeSpin adaptation

To further investigate the impact of C.NEST mixing, a second validation case was designed to determine if clones derived from static versus mixing workflows would differ in their adaptation to TubeSpin culture when simultaneously transferred from the 24-well stage (Figure 7).

**Figure 7.**
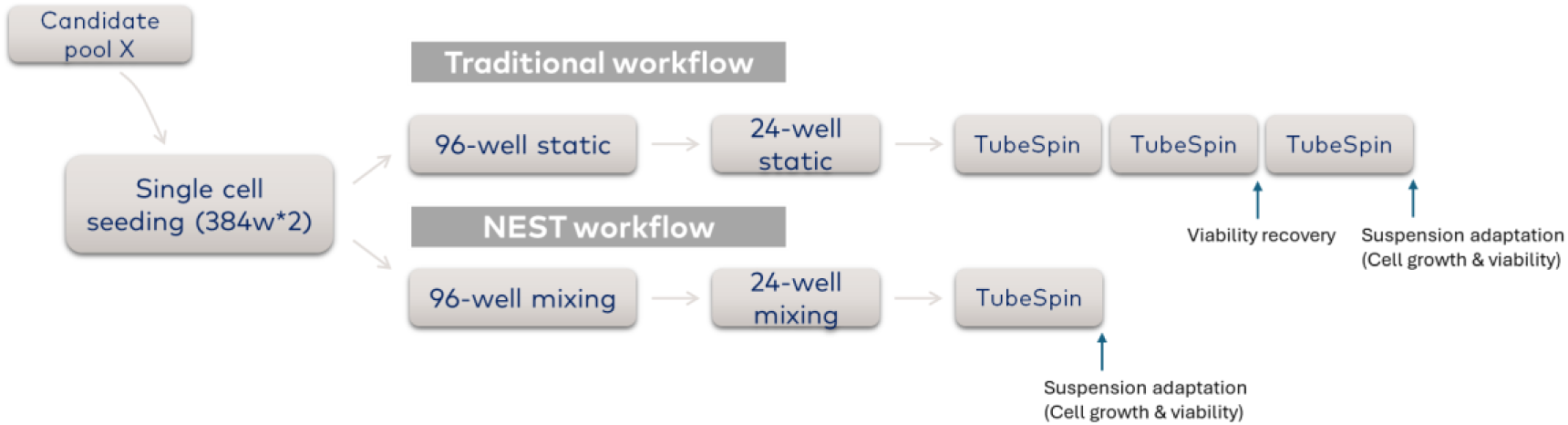
Case 2 experimental workflow. Schematic illustration of the clone expansion workflow under static and C.NEST mixing conditions. Clones derived from 384-well single-cell seeding were expanded through 96-well and 24-well culture, followed by TubeSpin adaptation. Confluency, VCD, and IgG titer were used as sequential selection criteria to evaluate clone performance at each stage.

From candidate pool X, single cells were isolated and deposited in 384-well plates. Confluency assessment on day 14 served as the first checkpoint, with wells exceeding 30% confluency eligible for transfer (Figure 8A). At the 96-well stage, clone selection was based directly on IgG titers rather than confluency due to imaging artifacts such as the ring effect under mixing conditions (Figure 8B). Twenty-four top-producing clones from both workflows were expanded into 24-well plates.

**Figure 8.**
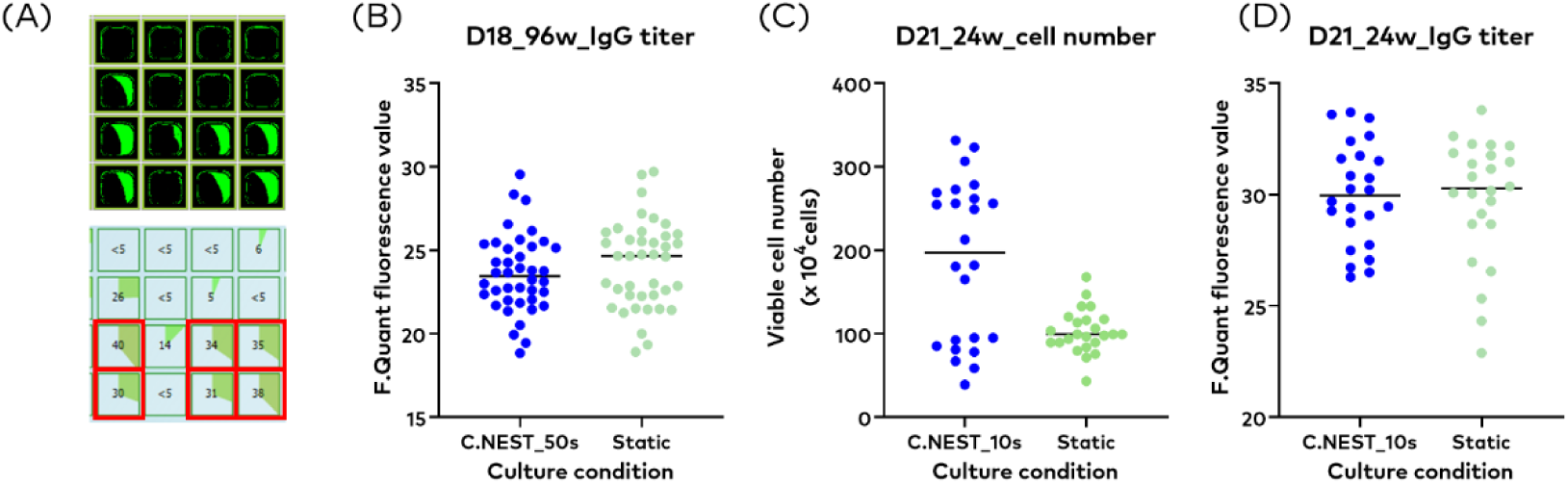
Case 2 –Clone selection and expansion from 384-well to 24-well. (A) Confluency assessment of 384-well single-cell culture used as the criterion for transferring clones into 96-well plates. Wells exceeding 30% confluency were selected for further expansion. Representative CloneSelect Imager images are shown.(B) IgG titer analysis at day 18 (D18) in 96-well plates served as the primary criterion for clone selection. Twenty-four high-titer clones from both static and mixing conditions were advanced to the 24-well stage.(C) VCD measured after 3 days of 24-well culture (D21). C.NEST-derived clones exhibited significantly higher viable cell numbers compared to static-derived clones.(D) IgG titer measured after 3 days of 24-well culture (D21). The top 8 performing clones were selected and transferred to the TubeSpin stage for scale-up under shaking culture.

Following transfer, clones pre-expanded with C.NEST mixing exhibited significantly higher viable cell density compared with static-derived clones, while IgG titers remained comparable (Figure 8C, 8D). These results indicate that C.NEST mixing improved cell expansion without compromising productivity during early-scale expansion.

On day 24, both static- and mixing-derived clones were transferred to TubeSpin bioreactors. After 3 days of TubeSpin culture, mixing-derived clones showed higher viable cell densities than static-derived clones (Figure 9A). By day 28, static-derived clones began to recover after the first passage at 3 × 10⁵ cells/mL, but their VCD remained lower than that of mixing-derived clones (Figure 9B). By day 31, static-derived clones achieved comparable cell densities after the second passage (Figure 9C). These findings demonstrate that pre-expansion under C.NEST mixing reduced the adaptation time required in TubeSpin culture.

**Figure 9.**
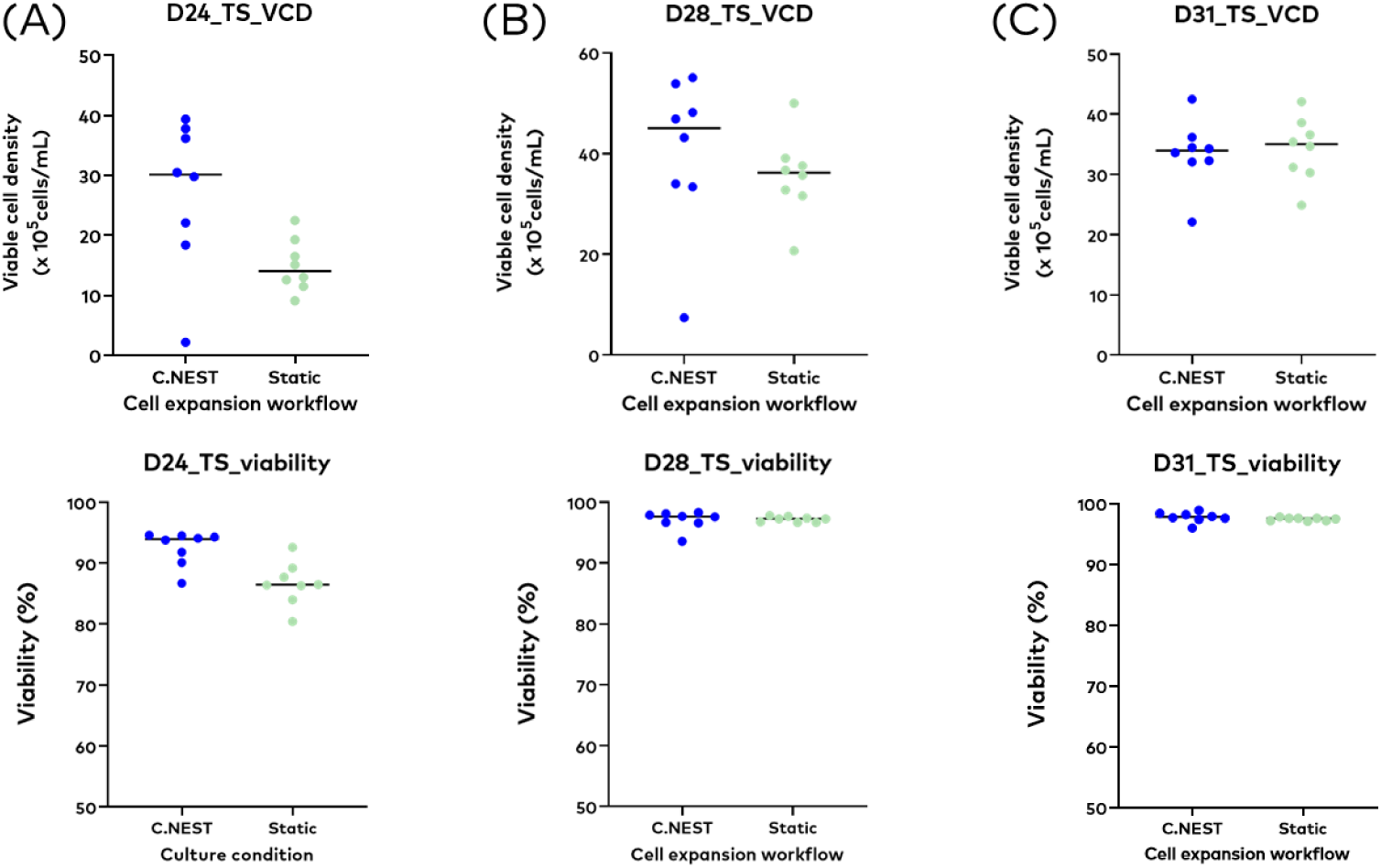
Case 2 – Clone performance in TubeSpin adaptation. (A) At day 24 (D24), after 3 days of TubeSpin culture, clones pre-expanded under C.NEST mixing showed significantly higher VCD compared to clones from static pre-culture.(B) At day 28 (D28), after the first passage at a seeding density of 3 × 10⁵ cells/mL, static-derived clones showed recovery in viability but still exhibited lower VCD than mixing-derived clones.(C) By day 31 (D31), after the second passage, static-derived clones reached comparable VCD levels to mixing-derived clones. Batch culture was then initiated to further evaluate clone performance.

To evaluate clone performance at production scale, top clones from both workflows were subjected to batch culture. Clones derived from mixing exhibited a wider distribution of viable cell densities and IgG titers, with outlier clones achieving markedly higher performance compared to static-derived clones (Figure 10). In contrast, static-derived clones showed narrower distributions and lacked standout high producers, suggesting that mixing culture may reveal superior clones otherwise overlooked in static workflows.

**Figure 10.**
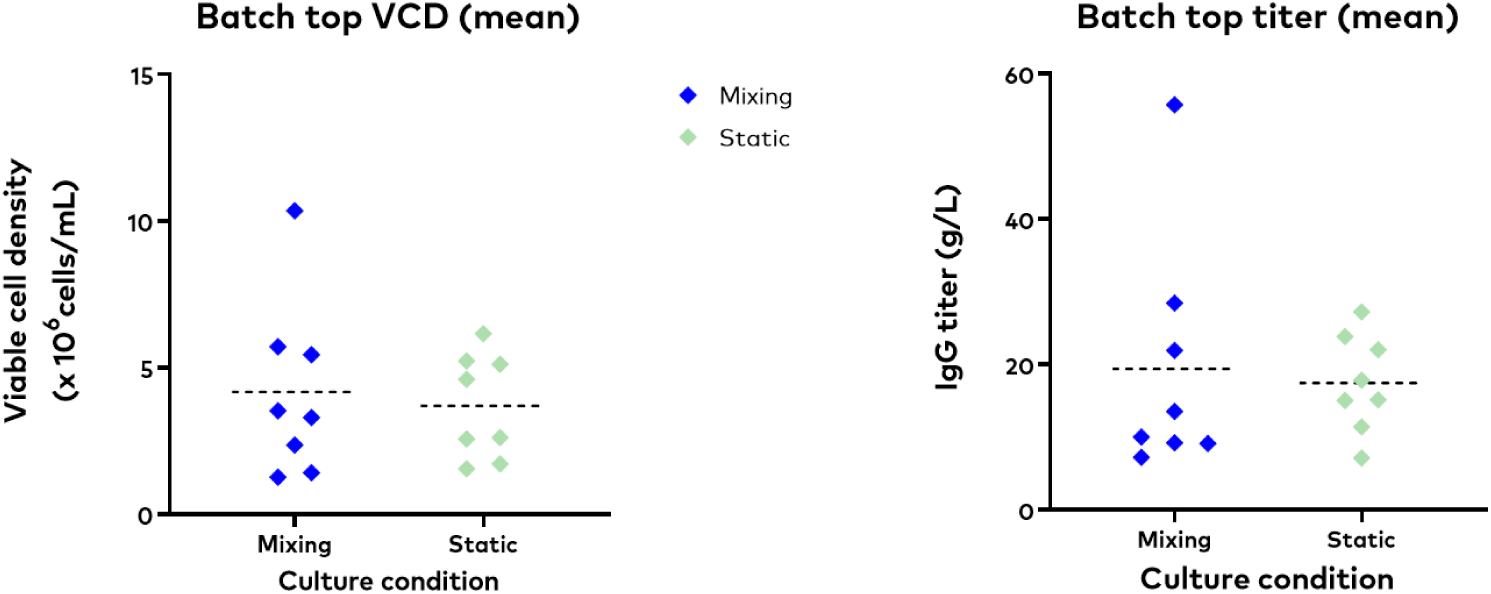
Case 2 –Batch culture performance of clones derived from mixing vs static workflows. Both workflows originated from the same clone pool.(A) Batch top VCD: In batch culture, mixing-derived clones displayed a wider distribution of VCD, with clear outliers showing much higher cell numbers. Static-derived clones, however, showed a narrower distribution without outlier performance.(B) Batch top IgG titer: Similarly, mixing-derived clones exhibited a broader range of titers with evident outliers, while static-derived clones maintained lower variation and lacked standout producers. Together, these results indicate that mixing culture can reveal outlier high-performing clones from the same clone pool, which are not captured by static culture.

The complete Case 2 workflow is summarized in Figure 11, illustrating the sequential checkpoints from 384-well SCC through microplate expansion and TubeSpin adaptation. Compared with the static workflow, C.NEST mixing facilitated earlier adaptation, accelerated clone expansion, and revealed outlier high-performing clones during batch evaluation. These results confirm that integrating C.NEST mixing into early clone selection not only reduces suspension adaptation time but also enhances the ability to identify top producers for scale-up.

**Figure 11.**
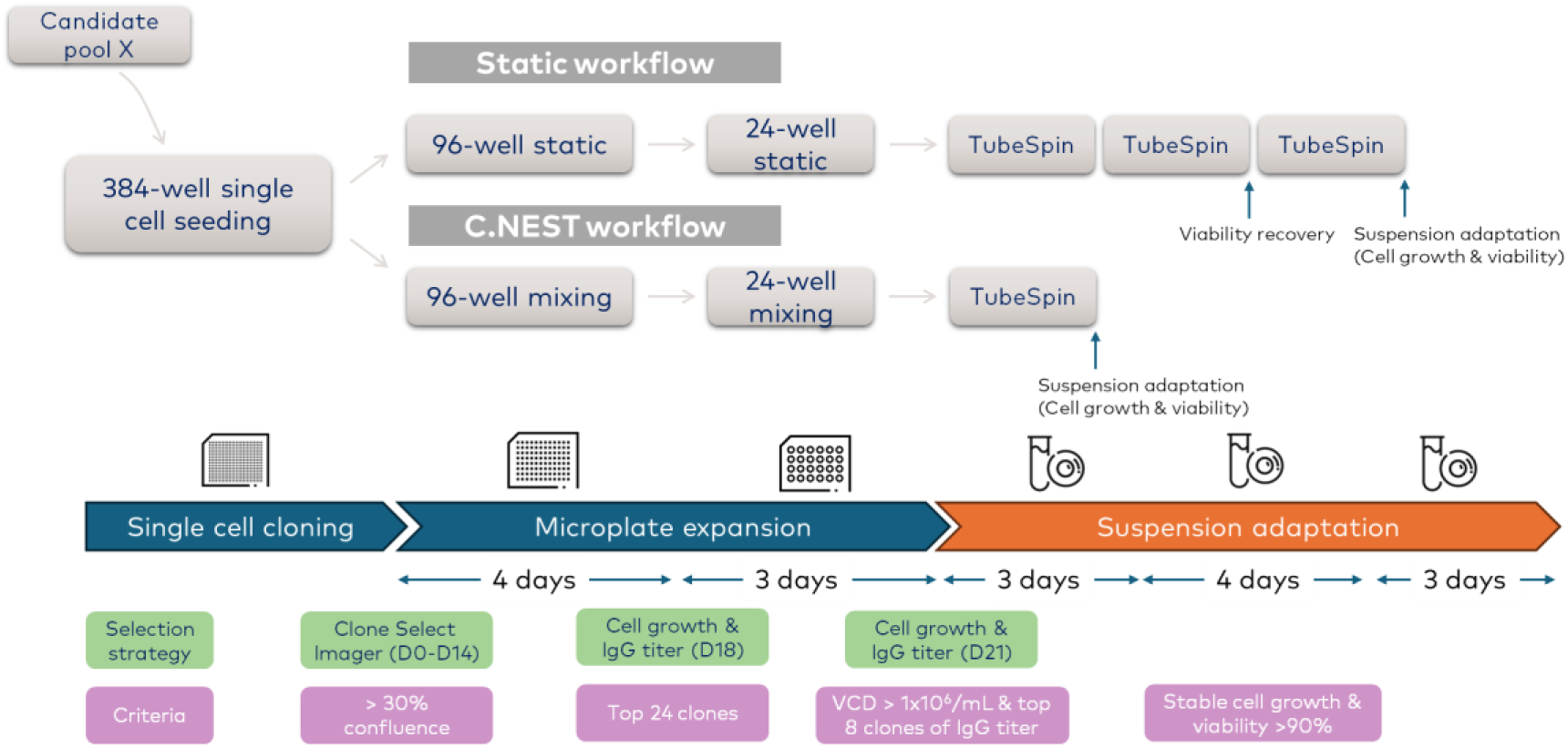
Case 2 workflow timeline for clone selection and expansion. Schematic overview of the static and C.NEST workflows starting from 384-well single-cell seeding (candidate pool X). **Static workflow**: Clones expanded sequentially through 96-well and 24-well static plates before transferring to TubeSpin, requiring additional passages for viability recovery and extended suspension adaptation. **C.NEST workflow**: Clones expanded under 96-well and 24-well mixing conditions, which enabled earlier suspension adaptation and direct transition to TubeSpin culture. The timeline (day 0–31) illustrates key selection checkpoints: **Day 14**: Confluency >30% in 384-well plates (CloneSelect Imager).Day 18: IgG titer measured in 96-well plates; top 24 clones advanced. **Day 21**: VCD >1 × 10⁶ cells/mL and IgG titer ranking in 24-well culture; top 8 clones selected. **Day 24–31**: TubeSpin expansion with suspension adaptation, confirming stable growth and >90% viability before batch culture evaluation.

Across both validation cases, C.NEST mixing consistently accelerated early clone expansion by facilitating suspension adaptation and increasing viable cell density compared with static workflows. This advantage was demonstrated in Case 1 during the 24-well to TubeSpin transition, and was further confirmed in Case 2, where synchronized TubeSpin transfer highlighted faster recovery and revealed outlier high-producing clones during batch evaluation.

### C.NEST-enabled acceleration of single cell cloning from mini-pool candidate

These findings corroborate C.NEST mixing as a robust strategy to shorten adaptation timelines and improve clone assessment; to demonstrate its practical utility in a full SCC workflow, we conducted a stepwise process beginning from candidate mini-pool (Figure 12). Mini-pool was first seeded into 384-well plates and expanded under static conditions. On day 10, wells that reached more than 10% confluency were transferred into 96-well mixing culture. At this stage, IgG titers were measured, and clones with values above 29.41 were selected for further expansion in 24-well mixing culture. After 3 days, VCD and IgG titers were quantified, and high-producers with titers greater than 32.98 were transferred to SF125 spin tubes or shake flasks for scale-up. Finally, batch culture was performed to validate clone performance, with both IgG titers and specific productivity (qP) measured as final readouts.

**Figure 12.**
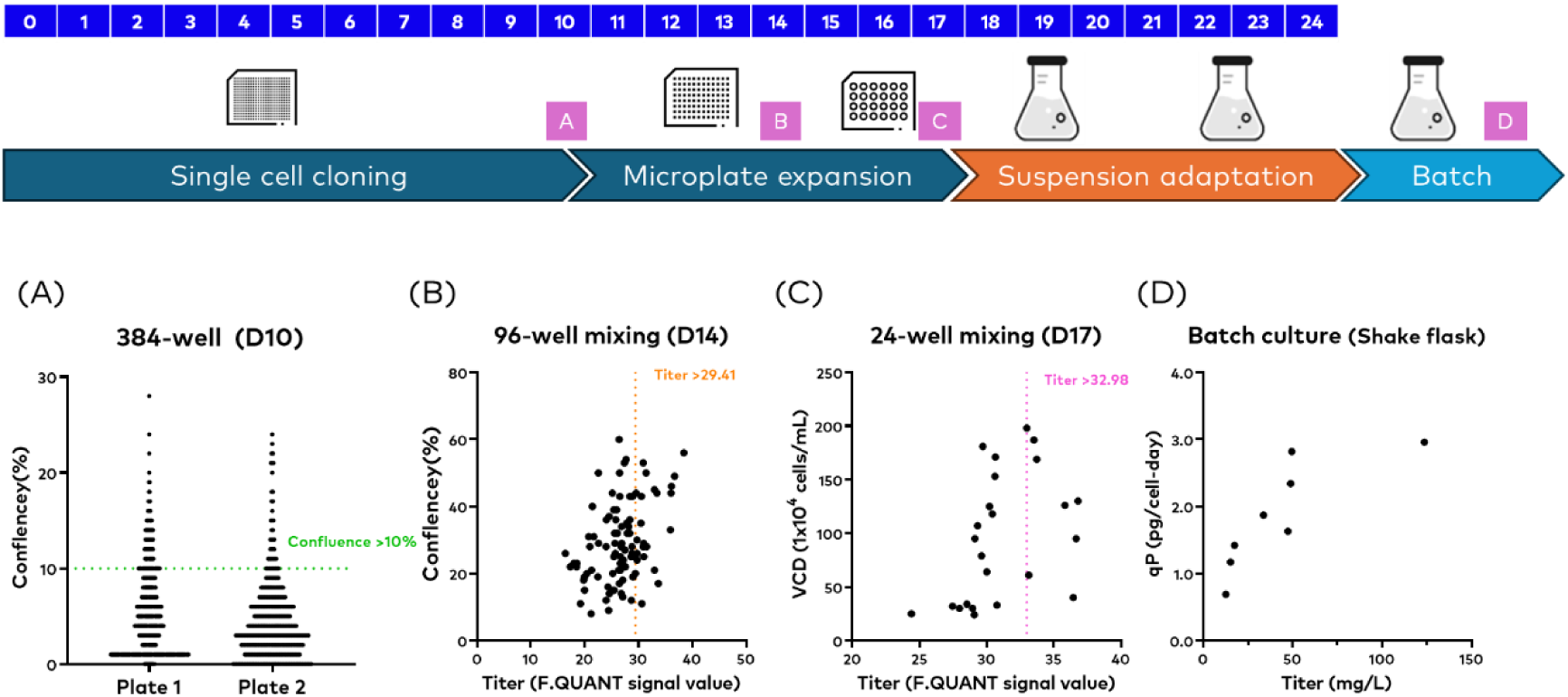
Application of C.NEST mixing in a single-cell cloning (SCC) workflow. (A) Single-cell cloning: Mini-pools were seeded into 384-well plates under static conditions. On day 10, wells with confluency greater than 10% were selected for transfer into 96-well mixing culture.(B) 96-well expansion: IgG titers were measured, and clones with titers above 29.41 were selected for further expansion.(C) 24-well mixing culture: After 3 days, VCD and IgG titers were quantified. Clones with titers greater than 32.98 were advanced into suspension culture (SF125 spin tubes or shake flasks).(D) Batch culture: Selected high-producing clones were validated for productivity, with both IgG titer and specific productivity (qP) measured to confirm consistent performance.

The results confirmed the effectiveness of this integrated workflow in enriching high-performing clones. On day 10 in 384-well plates, wells with confluency above 10% were selected for transfer (Figure 12A). At the 96-well stage, titer analysis identified clones with titers above 29.41 (Figure 12B). These clones were expanded into 24-well mixing culture, where both VCD and IgG titers were measured after 3 days, and clones exceeding a titer threshold of 32.98 were picked to the next stage (Figure 12C). Batch culture evaluation confirmed clone productivity, with qP plotted against IgG titers, demonstrating consistency of antibody production across selected clones (Figure 12D).

A final schematic (Figure 13) summarizes the integrated SCC workflow, highlighting how C.NEST mixing shortens the overall process by approximately one week compared to traditional static culture. This systematic approach enables efficient enrichment, expansion, and validation of high performing clones while maintaining robust suspension adaptation.

**Figure 13.**
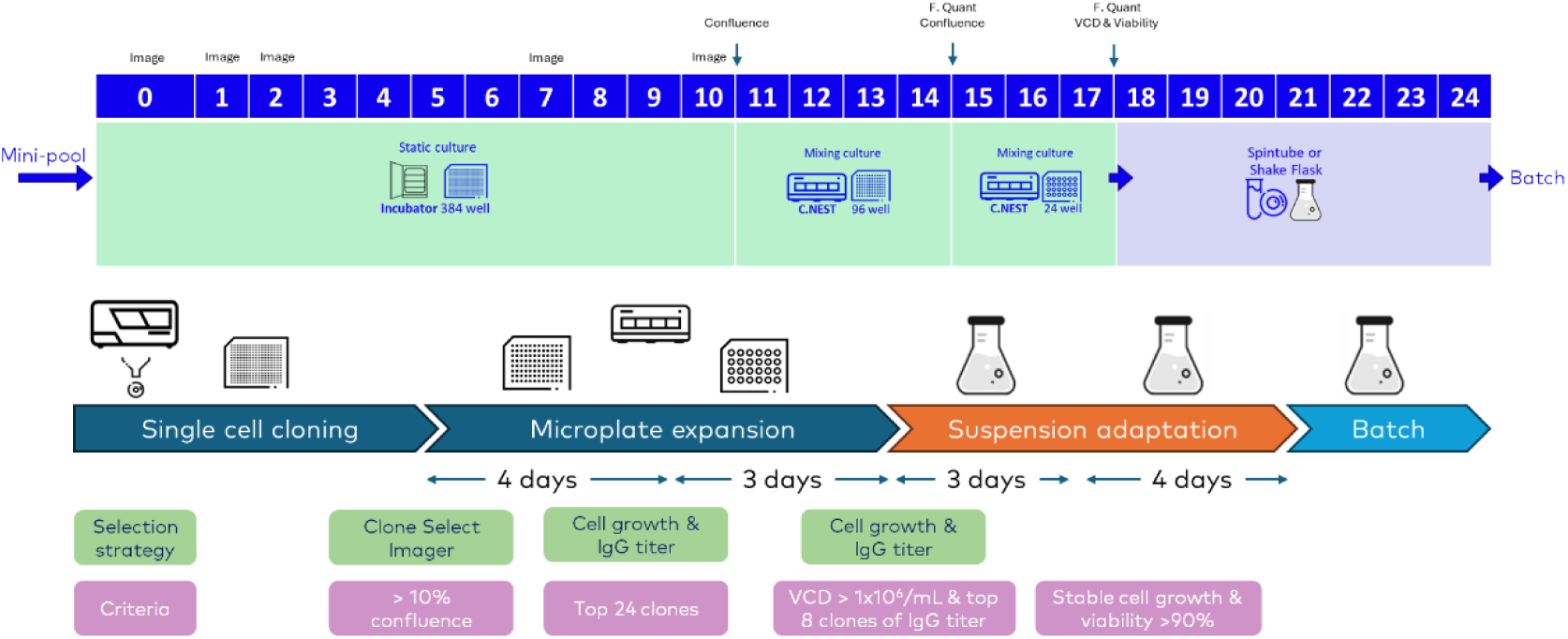
Summary of C.NEST-integrated single-cell cloning (SCC) workflow. Mini-pools were seeded into 384-well static culture, with wells exceeding 10% confluency by day 10 transferred to 96-well mixing culture. Following three days of expansion, IgG titer analysis enabled the selection of the top 24 clones, which were further expanded in 24-well mixing culture. After an additional three days, VCD and IgG titer were quantified, and the top eight clones were transferred to spin tube or shake flask culture for suspension adaptation. Stable cell growth and viability greater than 90% were achieved within four days, after which batch culture was performed for productivity validation. This workflow illustrates how C.NEST mixing enables earlier suspension adaptation and more reliable enrichment of high-performing clones, while reducing the overall timeline compared with static culture workflows.

## Discussion

The results of this study demonstrate the effectiveness of microplate agitation culture (C.NEST) in accelerating clone expansion and reducing suspension adaptation time during CHO-K1 cell line development. The transition from static to suspension in a new CLD campaign is traditionally a bottleneck phase in which cells often require extended recovery periods to achieve stable growth and viability (Wurm, 2013 [7]; Fischer et al., 2015 [8]). By introducing gentle mixing at the 96-well and 24-well stages, C.NEST enabled earlier suspension-like culture, which facilitated clone expansion and reduced adaptation delays when transferring to larger-scale shaking systems. Early-stage microplate cultures are often subject to local heterogeneity in dissolved oxygen, pH, and nutrient gradients, which can affect clone behavior and introduce bias in selection outcomes (Fisher et al., 2021 [9]; Neuss et al., 2025 [10]). The incorporation of controlled pneumatic mixing minimizes these gradients, improving mass transfer and ensuring more consistent microenvironments across wells.

Our findings highlight several key advantages of C.NEST’s pneumatic mixing. First, cell growth and viability were consistently higher in clones cultured under mixing compared with static conditions, supporting earlier expansion and scale-up. Second, the system improved clone selection accuracy by ensuring that differences in clone performance reflected inherent cellular characteristics rather than adaptation-related variability. This was evident in batch culture evaluation, where mixing-derived clones revealed broader productivity distributions and successfully isolated outlier high-producers that were missed by static culture. These results suggest that workflows lacking early mixing may underestimate the potential of certain clones, an observation consistent with previous reports that early culture conditions can influence long-term clone ranking and productivity outcomes (Wurm et al., 2017 [11]; Dorn et al., 2025 [12]).

The pneumatic-driven mixing mechanism employed in C.NEST represents a key distinction from conventional orbital or mechanical agitation systems. By alternating gas pressure and vacuum within the chamber, the platform generates a gentle, reciprocating convective flow that promotes uniform oxygen and nutrient distribution without subjecting cells to excessive mechanical shear. This mechanism likely contributes to the consistent improvements in viable cell density observed under mixing conditions. The enhanced homogeneity of the culture environment supports more stable cell proliferation and viability, addressing common limitations of static microplate systems where local gradients in oxygen and nutrient availability can constrain growth.

In the context of regulatory requirements, SCC remains the cornerstone for ensuring monoclonality and product consistency (ICH Q5D, 1998 [4]; EMA, 2006 [5]; FDA, 1992 [6]). However, the practical integration of C.NEST into both single-cell and mini-pool workflows demonstrates that microplate agitation systems can enhance not only operational efficiency but also data reliability for clone ranking. By shortening the suspension adaptation phase by approximately one week, the C.NEST platform can implement a fast-to-clinic/market strategy in biopharmaceutical manufacturing (Walsh, 2022 [1]). Furthermore, early adaptation conditions have been reported to influence the lag phase and robustness of suspension CHO cultures, supporting our conclusion that early-stage controlled mixing provides both physiological and operational advantages.

Furthermore, the principles demonstrated here align with ongoing efforts in CHO cell engineering and bioprocess optimization, where early control of growth conditions directly impacts productivity and stability (Kim et al., 2012 [2]; Fischer et al., 2015 [9]). The ability to integrate controlled mixing into standard CLD workflows provides a practical solution for improving clone recovery, minimizing variability, and enhancing throughput. Further investigation of this approach is warranted by combining microplate agitation with real-time metabolic monitoring or adaptive feeding strategies to optimize clone selection.

In conclusion, the unique mixing mode within C.NEST provides a robust platform for accelerating cell line development. By improving early-stage suspension adaptation and enabling more reliable clone performance assessment, this technology offers a streamlined pathway from SCC to batch validation, with the potential to reduce development timelines and improve the likelihood of identifying high-yield clones.

## Author Contributions

Shih-Pei Lin conceived and designed the study, performed the experiments, collected and analyzed the data, and wrote the manuscript. Ching-Nan Lin contributed to the study design, data analysis, and interpretation. Wei-Rou Wan assisted with experimental execution and data collection. Cheng-Han Tsai served as the corresponding author and was responsible for resource coordination and project supervision. All authors reviewed and approved the final version of the manuscript.

## Declaration of generative AI and AI-assisted technologies in the manuscript preparation process

During the preparation of this manuscript, the authors used ChatGPT, a generative AI tool, for assistance in manuscript drafting and English editing. The authors reviewed and revised all AI-assisted content and took full responsibility for the integrity, originality, and accuracy of the final published article.

